# Low oxygen arrests *Babesia duncani* schizonts and leads to increased drug susceptibilities in hamster erythrocytes

**DOI:** 10.1101/2021.01.11.426147

**Authors:** Yumin Zhang, Hector Alvarez-Manzo, Ying Zhang

## Abstract

We studied the effect of oxygen concentrations on the in vitro growth and drug susceptibility of *Babesia duncani*. We found that the growth of *B. duncani* required high level oxygen and the culture condition at ambient aerobic condition (21% O_2_) was optimal. Compared with ambient air, our results further showed that low oxygen (6-16%) could arrest *B. duncani* schizonts and lead to high susceptibilities to antiparasitic drugs atovaquone, pyrimethamine, quinine, and chloroquine at certain concentrations in vitro. Drug susceptibilities of other *Babesia* spp impacted by O_2_ levels need to be studied in the future, and this study indicates that culturing conditions of *Babesia* spp should be considered and reestablished for generating more comparable and reliable results in drug research in the future.

**Graphical abstract:** 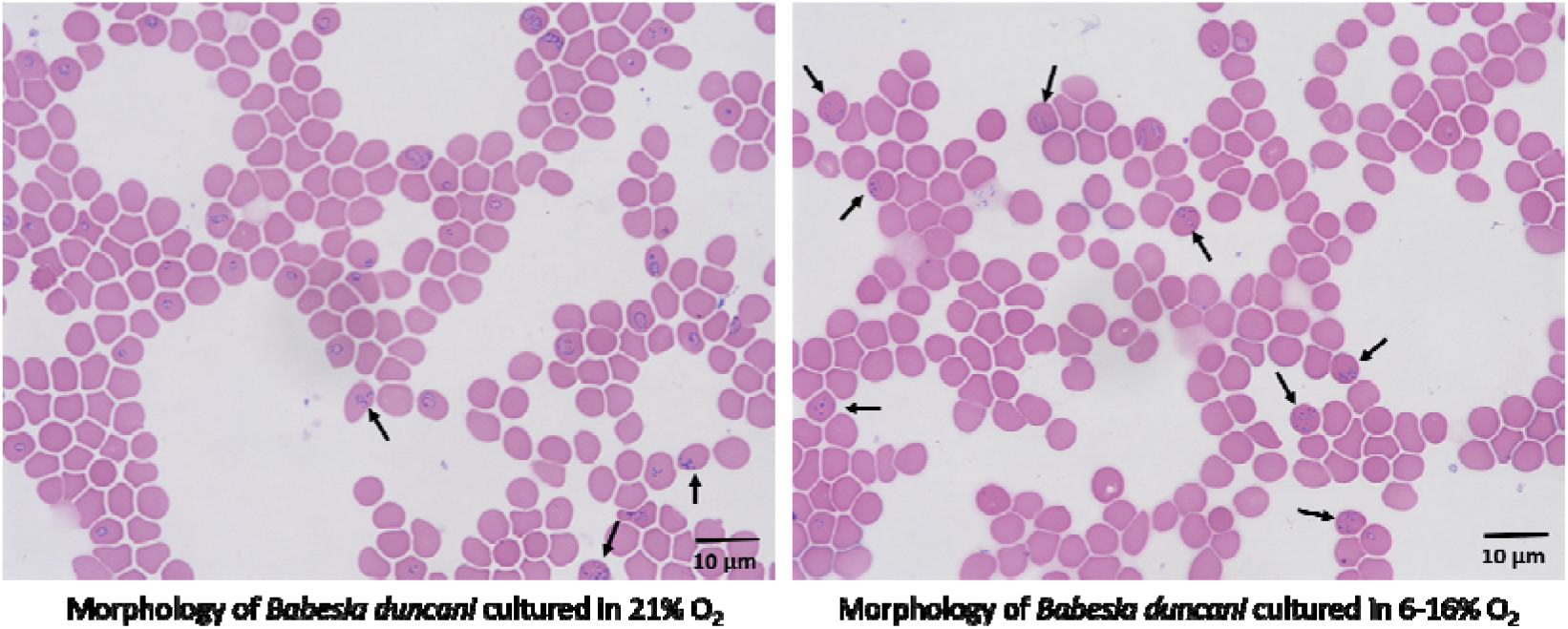

**Highlights:**

- The in vitro growth of *B. duncani* requires high level oxygen and the culture condition of 5% CO_2_ ambient air is optimal
- Low level oxygen results *B. duncani* in low growth rate and high schizont ratio in vitro
- Compared with 5% CO_2_ ambient air, in vitro drug susceptibilities of *B. duncani* can be significantly increased when cultured in microaerobic condition

---

Babesiosis is one of the most important tick-borne infectious diseases in Europe and North America and has been added to the list of nationally notifiable disease by the Centers for Disease Control and Prevention (CDC) of the United States. The clinical symptoms of babesiosis patients range from flu-like mild to malaria-like severe manifestations and can even cause death in elderly and immunocompromised patients. However, the current therapeutic regimen to human babesiosis is suboptimal with significant side effects and occasionally recrudescence (Kjemtrup and Conrad, 2000).

For development of more effective treatments, continuous in vitro culture of *Babesia* spp is one of the most critical techniques to explore active drugs. In the past decade, dozens of promising anti-babesia compounds have been identified with this technique. The first attempt to establish in vitro cultivation of *Babesia* spp in any real sense was in 1980, when Timms cultivated *B. bovis*, *B. bigemina*, and *B. rodhaini* in vitro up to 96 h using the candle jar technique which mimicked the in vitro culture of *Plasmodium falciparum* by Trager and Janson (Jensen and Trager, 1977; Timms, 1980). Subsequently, Levy and Ristic successfully carried out continuous cultivation of *B. bovis* in a stationary microaerophilic phase where oxygen had decreased to the level that could darken red blood cells laid on plate bottom (Levy and Ristic, 1980). In 1985, Vega *et al.* further demonstrated that high oxygen (5.6% - normal air) is detrimental to in vitro growth of *B. bigemina* (Vega et al., 1985). Thus, it seems that *Babesia* spp are microaerophilic organisms like *P. falciparum*. However, the in vitro cultivations of two important human babesiosis causative agents *B. divergens* and *B. duncani* were set in different O_2_ levels in some studies for drug activity testing or drug screening (Abraham et al., 2018; Brasseur et al., 1998; Nehrbass-Stuedli et al., 2011; Zhang et al., 2020), which have could lead to divergent results in anti-malarial drug studies (Branco et al., 2018; Duffy and Avery, 2018). Thus, two interesting questions are raised here: first, does *B. duncani* grow well in low O_2_; second, is O_2_ level a factor that affects the anti-babesia drug activity in vitro?

To address these questions, here we studied the effects of O_2_ levels on the growth and drug tolerance of *B. duncani* (WA1 strain), the latest human babesiosis pathogen to be identified. In continuous culture model in hamster erythrocytes cultivated in 5% CO_2_ ambient air with humidity (Zhang et al., 2020), we found *B. duncani* schizont was the major morphology in old cultures (5-6 culture days, 0.5% of initial parasitemia) and this morphology rarely appeared in young cultures (Figure 1A). These schizonts appearing as budding four daughter merozoites in erythrocytes and displayed several different scattered distributions (Figure 1, B1-B6). Considering culture medium changed daily and erythrocytes turned dark in old cultures, we hypothesize that low oxygen environment resulting from rapid oxygen consumption by parasites in high parasitemia is related to the increasing percentage of schizonts. To confirm this, when parasitemia reached to 1.5-2% cultivated in 5% CO_2_ atmosphere, we split the *B. duncani* cultures into anaerobic, microaerobic, and aerobic conditions without other culturing condition change to evaluate the effects of oxygen concentrations on parasite growth and morphologic development. In this study, anaerobic (0.7-1% O_2_) and microaerobic (6-16% O_2_) conditions were generated by BD GasPak EZ container systems with CampyPak container system sachets and anaerobe container system sachets, respectively, and 5% CO_2_ atmosphere was viewed as aerobic condition (21% O_2_, approximately). Even though the final CO_2_ concentrations in BD GasPak EZ container systems would be various from 2% to 13% according to the manufactory’s instructions, our results indicated that the rate of growth and morphologic development of *B. duncani* did not show significant difference among 2, 5, and 13% CO_2_ atmosphere air the cultures incubated in, but *B. duncani* did not grow in atmosphere air deprived CO_2_ (data not shown). Culture medium covered erythrocytes replaced every day with fresh medium which had been co-incubated in respective culture environments to minimize the impact of the fluctuation of nutrition and dissolving oxygen.

**Figure 1.**
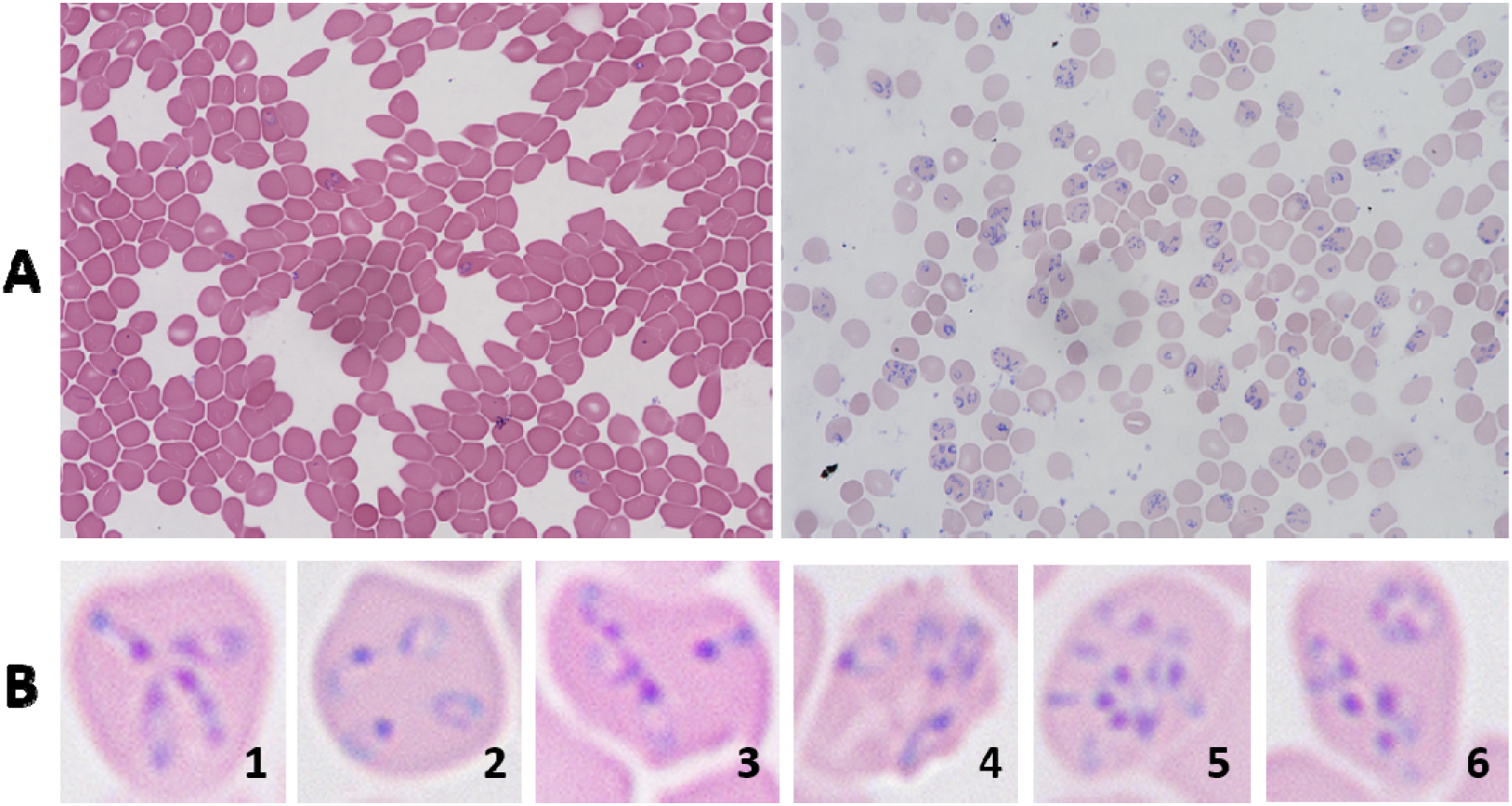
(A) *B. duncani*-infected hamster erythrocytes observed at 1,000× magnification in young (2 days old; left) and old (5-6 days old; right) cultures with parasitemia of approximate 1.5% and 40%, respectively. The growth of parasites started from 0.5% parasitemia; (B) *B. duncani* schizonts appearing as budding four daughter merozoites in erythrocytes displayed several different scattered distributions. B1, Maltese Cross tetrad; B2, four daughter merozoites connected in pairs; B3, three daughter merozoites assembled with one in isolation; B4, the moment of matured merozoites bursting erythrocyte; B5, two schizonts in one erythrocyte; B6, a schizont accompanied with ring form trophozoite;

Our results showed that the growth of *B. duncani* incubated in microaerophilic condition reached to peak parasitemia of approximate 16% at day 5. As a comparison, the parasitemia of cultures incubated in aerobic condition could finally reach to approximate 40% at day 5 before erythrocyte lysis, and it grew faster than that in microaerophilic condition after 1 day (Figure 2). Growth of *B. duncani* stopped when incubated in anaerobic condition, and no infected erythrocyte was observed starting at day 3 (Figure 2). This indicates that *B. duncani* is an oxygen preferred organism, and the growth of *B. duncani* requires more oxygen than only 2-5% O_2_ employed in previous studies which may not be the optimal in vitro growth condition for *B. duncani* (Abraham et al., 2018; McCormack et al., 2019). It can partly explain why the *B. duncani* cultures obtained in ambient air was able to reach to much higher parasitemia than reported parasitemias, i.e., 40% in our study vs 20% or less previously (Abraham et al., 2018; McCormack et al., 2019). Of note, the pericellular O_2_ around the parasites and erythrocytes laid on plate bottom must be lower than environmental O_2_ due to the culture medium with depth of 0.7cm in our culturing system which acted as an oxygen barrier.

**Figure 2.**
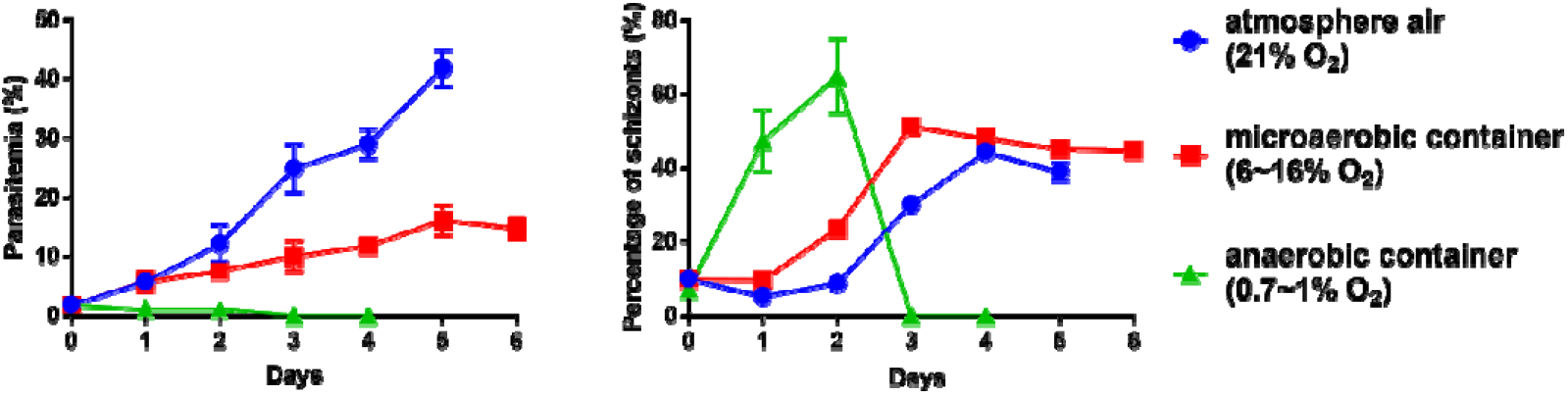
Growth (left) and percentage of schizonts (right) of *B. duncani* cultures incubated in aerobic, microaerobic, and anaerobic environments;

In the human body, mature *P. falciparum* parasites (trophozoites and schizonts) exhibit a preference for the venules where relatively high O_2_ levels are present (Milner et al., 2015), Research on the effect of oxygen on the growth of *Babesia* spp has not been reported in vitro or in vivo. In contrast, our in vitro results showed *B. duncani* schizonts were more likely arrested in anaerobic and microaerobic conditions than ambient oxygen rich condition (Figure 2). Specifically, even though *B. duncani* did not grow in anaerobic container (0.7-1% O_2_), the ratio of schizonts in infected erythrocytes soared immediately at the beginning and could finally reach to nearly 70% schizonts before all parasites died out (Figure 2). The ratios of schizonts were increasing along with increasing parasitemia and oxygen consumption in both microaerobic and aerobic conditions, but the ratios of schizonts cultured in microaerobic condition were higher than that cultured in aerobic condition at each day and entered plateau one day earlier. This indicates that low O_2_ is able to extend the duration of *B. duncani* schizonts in vitro or prevent schizonts from lysing erythrocytes to generate merozoites, which could also happen in late culture phase. In contrast, Briolant et al. found that *P. falciparum* clones 3D7 and W2 had higher percentages of schizonts when inoculated in 21% O_2_ than that inoculated in 5% O_2_, even though the growth of these two *P. falciparum* clones did not show significant difference in 21% and 5% O_2_ (Briolant et al., 2007).

Slight variations of in vitro culture environments have demonstrated significant impact on the conclusions of *P. falciparum* drug resistance (Duffy and Avery, 2018). In our study, we found that *B. duncani* cultured in microaerobic condition showed significantly higher susceptibility to four anti-parasite drugs atovaquone, pyrimethamine, quinine, and chloroquine at 0.25-50 μM than that cultured in aerobic condition, except that atovaquone had almost 100% inhibitory effects against *B. duncani* at 2.5, 25, and 50 μM in both aerobic and microaerobic conditions (Figure 3). The most variation of inhibitory effects occurred in chloroquine exposure, which had 65-100% inhibitory effect on *B. duncani* at 0.25 and 2.5 μM in microaerobic condition, but the effect was only 9-34% at the same drug concentrations in aerobic condition (Figure 3). As mentioned above, given *P. falciparum* 3D7 and W2 have the opposite manifestation of ratios of schizonts in different O_2_ levels compared to *B. duncani* in this study, it is not surprising that *P. falciparum* 3D7 and W2 were more susceptible to chloroquine in aerobic condition than that in microaerobic condition (Briolant et al., 2007), which is dissimilar to the chloroquine susceptibility of *B. duncani* as shown in our study. We speculated that these different consequences could result from the same reason that schizonts are more susceptible to chloroquine due to chloroquine accumulation might be much greater in schizont stage than in ring stage (Yayon et al., 1983). Besides, some compounds have shown various activities against highly synchronous age defined *P. falciparum* parasites (Duffy and Avery, 2017), however, to date there is no way to study drug susceptibility of *Babesia* spp at a certain life cycle stage due to the lack of effective synchronization of *Babesia* spp. Drug susceptibilities of other *Babesia* spp impacted by O_2_ levels need to be studied in the future, and culturing conditions of *Babesia* spp should be reestablished for generating more comparable and reliable results in drug research and the relevant conditions that reflect in vivo drug treatment effect should be determined to identify more effective drugs.

**Figure 3.**
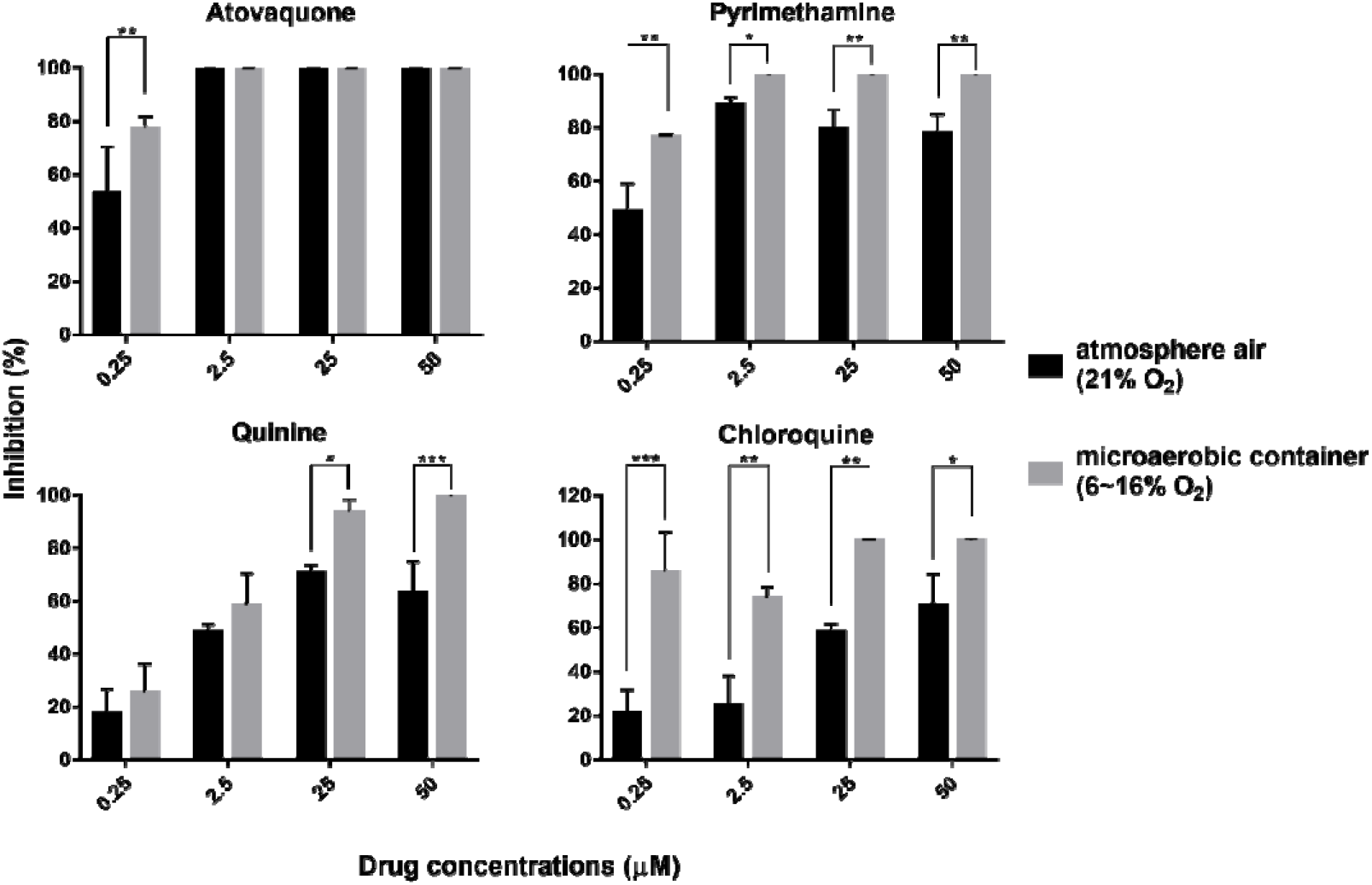
*B. duncani* incubated in aerobic and microaerobic conditions with a starting 1.5% parasitemia and 2.5% hematocrit (2 days old culture) treated with atovaquone, pyrimethamine, quinine, and chloroquine at 0.25, 2.5, 25, 50 μM after 3 days exposure.

